# Dissecting the first phased dikaryotic genomes of the wheat rust pathogen *Puccinia triticina* reveals the mechanisms of somatic exchange in nature

**DOI:** 10.1101/705475

**Authors:** Jing Qin Wu, Chongmei Dong, Long Song, Christina A. Cuomo, Robert F. Park

## Abstract

Although somatic hybridization (SH) has been proposed as a means of accelerating rust pathogen virulence evolution in the absence of sexual recombination, previous studies are limited to the laboratory and none have revealed how this process happens. Using long-read sequencing, we generated dikaryotic phased genomes and annotations for three Australian field-collected isolates of the wheat leaf rust pathogen (*Puccinia triticina; Pt*), including a putative asexual hybrid (Pt64) and two putative parental isolates (Pt104 and Pt53; 132-141 Mb,155-176 contigs, N_50_ of 1.9-2.1 Mb). The genetic dissection based on the high-quality phased genomes including whole-genome alignments, phylogenetic and syntenic analyses along with short-read sequencing of 27 additional *Pt* isolates convergently demonstrated that Pt64, which rendered several commercial hybrid wheat cultivars susceptible to leaf rust, arose from SH between isolates within the Pt53 and Pt104 lineages. Parentage analysis demonstrated the role of mitotic crossover in the derivation of both nuclei of Pt64. Within HD mating type genes, the distinct specificity regions in Pt64 and the distinct phylogenetic pattern of the remaining admixed isolates suggested high genetic variation in specificity-related regions on the *b* locus intrinsically associated with the SH. This study not only provided a fundamental platform for investigating genomic variation underlying virulence evolution in one of the most devastating wheat pathogens, but also offered an in-depth understanding of the mechanisms of naturally occurring SH. This asexual mechanism can be broadly exploited by any dikaryotic pathogen to accelerate virulence evolution, and understanding this process is both urgent and crucial for sustainable pathogen control.

**Importance:** Strategies to manage plant rust pathogens are challenged by the constant emergence of new virulence. Although somatic hybridization has been proposed as a means by which rusts could overcome host resistance rapidly and cause crop loss, there is very little evidence of this process in nature and the mechanisms underlying it are not known. This study generated and analysed the first dikaryotic phased genomes of the wheat leaf rust pathogen, identifying an isolate as a hybrid and for the first time unveiling parasexuality via mitotic crossover in a rust pathogen. The erosion of the resistance of several hybrid wheat cultivars in agriculture by the hybrid rust has important implications for breeding efforts targeting durable resistance and sustained rust control.

## Introduction

Leaf rust, caused by *Puccinia triticina* (*Pt*), is one of the most devastating diseases of wheat, affecting production in nearly all wheat growing regions worldwide and with annual yield losses reaching 40% (1). Because of its widespread and common occurrence, leaf rust is considered to be the most damaging of the three rust diseases of wheat. *Pt* has a complex life cycle including sexual and asexual stages, which is truncated in some parts of the world where sexual recombination does not occur due to the absence of the alternate host. The absence of sexual recombination in *Pt* in Australia and its geographic isolation from other wheat growing regions of the world have permitted continental-scale studies of the evolution of genetic variation in a large asexually reproducing fungal pathogen population, through the detection of phenotypes with new pathogenicity traits and combinations of rare pathogenic features (2). Such studies over much of the past 95 years have implicated mutation and periodic exotic incursion in the generation of genetic variation in *Pt* and several other cereal rust pathogens. Evidence for somatic hybridization (SH) in nature has only been reported twice in a cereal rust pathogen, in the wheat stem rust pathogen *P. graminis* f. sp. *tritici* (*Pgt*) (3), and in *Pt*. In *Pt*, pathotype 64-(6),(7),(10),11 (Pt64), first detected in Australia in 1990, was postulated to have arisen via hybridisation between two founding and genetically distinct pathotypes, 53-1,(6),(7),10,11 (Pt53; first detected in Australia in 1984) and 104-2,3,(6),(7),11 (Pt104; first detected in 1984) (4). Significantly, the putative hybrid Pt64 rendered several commercial hybrid wheat cultivars grown in north eastern Australia susceptible to leaf rust.

The mechanisms underlying SH in rust fungi are not known. Previous studies under laboratory conditions have suggested that it involves the anastomosis of hyphae followed by reassortment of whole nuclei and/or the parasexual cycle (nuclear fusion, chromosomal exchange, mitotic crossing over, and segregation) (5–7). While the exchange of partner nuclei between genotypes with neither nucleus in common is expected to generate no more than two recombinant types (e.g. in the rust fungi *Melampsora lini* and *Pst* (6, 7)), parasexuality similar to that described for *Aspergillus nidulans* was suggested based on the recovery of more than two recombinant pathogenicity types in *Pgt* and the oat crown rust pathogen *Puccinia coronata* f. sp. *avenae* (*Pca*) in laboratory studies (8). It remains unclear whether SH occurs in rust pathogens in nature, and the mechanisms underlying such a process in these fungi are unknown. Previous studies in *Ustilago maydis* suggested that incompatibility of mating type genes may preclude the formation of heterokaryons between vegetative cells (9). In the phylum Basidiomycota, two sets of mating type genes encoding for lipopeptide pheromones along with their cognate receptors and homeodomain transcription factors (HD1 and HD2) are essential for sexual reproduction (10). All of these genes were detected in a draft genome of *Pt* Race 1 (11). Furthermore, cytological studies on laboratory germinated urediniospores of *Pt* revealed germ tube fusion leading to hyphal anastomosis, demonstrating the possibility of asexual fusion between genetically different individuals with different virulence phenotypes and suggesting another means of combining different virulences and creating a heterokaryon with a broader virulence spectrum than the original isolates (12).

With the advent of next generation sequencing (NGS) technology, the draft genomes of wheat rust fungi including *Pt*, *Pgt*, and *P. striiformis* f. sp. *tritici* (*Pst*) have become available and several resequencing projects undertaken for effector mining. These projects include five *Pgt* isolates (13), 10 *Pst* isolates (14, 15), and 20 Australian *Pt* isolates comprising 10 pairs each differing in avirulence/virulence only to *Lr20* (16). While these resequencing studies have identified a small list of effector candidates for further validation, a more recent comparative study between an isolate of *Pgt* (Pgt279) and a derivative (Pgt632) isolate with virulence for *Sr50* successfully identified *AvrSr50* (17). Recently, third generation long-read sequencing has been used to assemble the first haplotype-phased reference genomes for the dikaryotic rust fungi *Pst* and *Pca*, which produced reference genomes with much higher contiguity than those based on short-read sequencing and uncovered novel genome features in a dikaryotic fashion (18, 19). In the present study, long-read sequencing technology was used to generate three haplotype-phased genome assemblies for the putative hybrid (Pt64) and parental isolates (Pt104 and Pt53), which are the first phased dikaryotic *Pt* genomes with the most complete information in terms of contiguity, haplotype-phased information, and gene content. By dissecting these high-quality genomes, the evolutionary trajectories of a rust SH in nature were revealed for the first time. In addition to providing solid evidence of SH in a dikaryotic pathogen in nature, our study unveiled an underlying mechanism of SH in a dikaryotic fungus. More complex than simple nuclear exchange as previously believed, this genetic process involved mitotic crossover in both nuclei of the hybrid isolate and all four parental nuclei contributed. Further analysis on HD mating types genes suggested that high genetic variation in specificity-related regions between HD genes on the *b* locus could be intrinsically associated with SH. The mechanism identified could be used by any dikaryotic pathogen in nature to evolve more rapidly to overcome host resistance in the absence of sexual recombination. Understanding this process will undoubtedly facilitate the development of more sustainable pathogen control strategies.

## Results and discussion

### Assembly and annotation of high quality, haplotype phased *Pt* genomes

We generated the first haplotype phased genomes for the wheat leaf rust pathogen *Puccinia triticina*, targeting three field collected isolates: a putative asexual hybrid (Pt64) and its two putative parental isolates (Pt104 and Pt53). These three isolates were sequenced using the PacBio Sequel system (average read length > 10 kb, 3-4 SMRT cells/sample, about 100-200 coverage) and Illumina (> 70 coverage), along with sequencing with Illumina of an additional 27 *Pt* isolates covering a wide range of pathotypes. The genomes were assembled by a Falcon, Falcon-Unzip and Quiver pipeline, followed by polishing and curation. The final curated *Pt* assemblies comprised 155-176 primary contigs (N_50_ of 1.9-2.1 Mb) and 531-843 haplotigs, totaling 132-141 Mb and 123-132 Mb, respectively (**Table 1**). They were approximately 92% complete, and only 3.7% fragmented BUSCO-predicted genes. When primary contigs and haplotigs were combined, missing BUSCO genes were as low as 2.4-2.6% (**Table 1**). The total interspersed repeats covered 58.2-59.8% in primary contigs and 56-57% in haplotigs using both *de novo* predicted repeats and fungal elements from RepBase (20) (**Table 1; supplementary file 1**). The total number of unique telomeres identified in the three isolates ranged from 10-17, which was in the range of previous estimates (12 telomere ends out of 6 haploid chromosomes) (21).

**Table 1.**
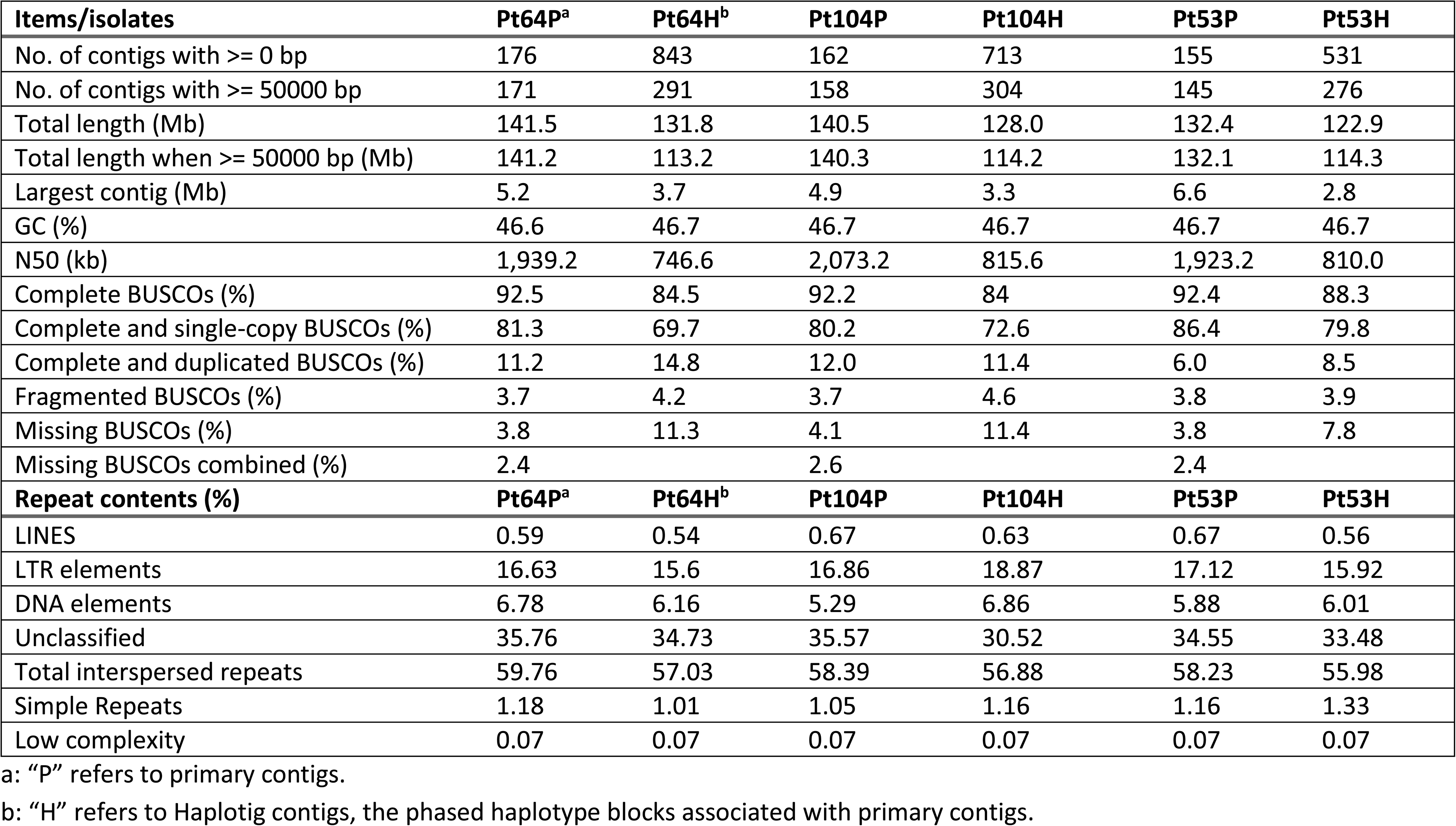
Assembly statistics, completeness evaluation and repeated sequence content in the phased genomes of 3 Pt isolates.

| Items/isolates | Pt64P<sup>a</sup> | Pt64H<sup>b</sup> | Pt104P | Pt104H | Pt53P | Pt53H |
| --- | --- | --- | --- | --- | --- | --- |
| No. of contigs with >= 0 bp | 176 | 843 | 162 | 713 | 155 | 531 |
| No. of contigs with >= 50000 bp | 171 | 291 | 158 | 304 | 145 | 276 |
| Total length (Mb) | 141.5 | 131.8 | 140.5 | 128.0 | 132.4 | 122.9 |
| Total length when >= 50000 bp (Mb) | 141.2 | 113.2 | 140.3 | 114.2 | 132.1 | 114.3 |
| Largest contig (Mb) | 5.2 | 3.7 | 4.9 | 3.3 | 6.6 | 2.8 |
| GC (%) | 46.6 | 46.7 | 46.7 | 46.7 | 46.7 | 46.7 |
| N50 (kb) | 1,939.2 | 746.6 | 2,073.2 | 815.6 | 1,923.2 | 810.0 |
| Complete BUSCOs (%) | 92.5 | 84.5 | 92.2 | 84 | 92.4 | 88.3 |
| Complete and single-copy BUSCOs (%) | 81.3 | 69.7 | 80.2 | 72.6 | 86.4 | 79.8 |
| Complete and duplicated BUSCOs (%) | 11.2 | 14.8 | 12.0 | 11.4 | 6.0 | 8.5 |
| Fragmented BUSCOs (%) | 3.7 | 4.2 | 3.7 | 4.6 | 3.8 | 3.9 |
| Missing BUSCOs (%) | 3.8 | 11.3 | 4.1 | 11.4 | 3.8 | 7.8 |
| Missing BUSCOs combined (%) | 2.4 |  | 2.6 |  | 2.4 |  |
| **Repeat contents (%)** | **Pt64P<sup>a</sup>** | **Pt64H<sup>b</sup>** | **Pt104P** | **Pt104H** | **Pt53P** | **Pt53H** |
| LINES | 0.59 | 0.54 | 0.67 | 0.63 | 0.67 | 0.56 |
| LTR elements | 16.63 | 15.6 | 16.86 | 18.87 | 17.12 | 15.92 |
| DNA elements | 6.78 | 6.16 | 5.29 | 6.86 | 5.88 | 6.01 |
| Unclassified | 35.76 | 34.73 | 35.57 | 30.52 | 34.55 | 33.48 |
| Total interspersed repeats | 59.76 | 57.03 | 58.39 | 56.88 | 58.23 | 55.98 |
| Simple Repeats | 1.18 | 1.01 | 1.05 | 1.16 | 1.16 | 1.33 |
| Low complexity | 0.07 | 0.07 | 0.07 | 0.07 | 0.07 | 0.07 |

a: "P" refers to primary contigs.
b: "H" refers to Haplotig contigs, the phased haplotype blocks associated with primary contigs.

For genome annotation of the three *Pt* isolates, the Funannotate v0.7.2 pipeline combining multiple lines of evidence such as RNA-seq of Pt64 at multiple time points, the proteome from *Pt* Race 1 (11), *ab initio* predicted transcript models, and ESTs from various life cycle stages of *Pt* (22), were used. This predicted 27,000-29,000 gene models for the primary contigs of each isolate and 26,000-28,000 for the haplotigs (**Table 2; Supplementary file 2**). While our predicted gene number is higher than that for *Pt* Race 1 (14,000-15,000) (11), it is very similar to another *Pt* study on races 77 and 106 (26,000-27,000) (23). The differences may be largely attributed to the differences in the isolates from different countries and in gene annotation and filtering methods, and keeping a relatively broader range of predicted genes may facilitate future efforts to identify effector protein encoding genes. Paired orthologs between the primary contigs and haplotigs within each isolate were identified by Proteinortho v5.16 (synteny mode). A total of 15,000-16,500 ortholog pairs were identified within each isolate, which were further classified into synteny allele pairs (about 10,000-12,500), chromosome allele pairs (about 2,500-3,300), and location-unmatched ortholog pairs (about 1,300-2,100; **Table 2; Supplementary file 3**) based on their location in relation to synteny blocks as described in the Methods section.

**Table 2.** Genes, allele pairs, and effectors in the phased-genomes of 3 Pt isolates.

|  | **Pt64P<sup>a</sup>** | **Pt64H<sup>b</sup>** | **Pt104P** | **Pt104H** | **Pt53P** | **Pt53H** |
| --- | --- | --- | --- | --- | --- | --- |
| **Gene and protein stats** |  |  |  |  |  |  |
| Total number of genes | 28,500 | 28,068 | 29,043 | 27,222 | 27,236 | 26,205 |
| Mean gene length | 1,380 | 1,354 | 1,378 | 1,350 | 1,386 | 1,365 |
| genome % covered by genes | 27.8 | 28.9 | 28.5 | 28.7 | 28.5 | 29.1 |
| Total number of proteins | 27,356 | 27,013 | 28,008 | 26,236 | 26,174 | 25,277 |
| **ProteinOrtho analysis across 6 genomes (3 primary contigs and 3 haplotigs)** |  |  |  |  |  |  |
| Pr. with at least 1 ortholog | 24,599 | 23,722 | 24,726 | 22,846 | 23,041 | 22,426 |
| Single copy orthologs<sup>c</sup> | 7,869 | 7,869 | 7,869 | 7,869 | 7,869 | 7,869 |
| Singletons<sup>d</sup> | 2,757 | 3,291 | 3,282 | 3,390 | 3,133 | 2,851 |
| **ProteinOrtho analysis within an isolate between primary contigs and haplotigs** |  |  |  |  |  |  |
| Synteny allele pairs<sup>e</sup> | 9966 | 9966 | 11331 | 11331 | 12462 | 12462 |
| Chromosome allele pairs<sup>f</sup> | 3274 | 3274 | 3021 | 3021 | 2569 | 2569 |
| unmatched ortholog pairs<sup>g</sup> | 2047 | 2047 | 1709 | 1709 | 1365 | 1365 |
| **Secretome stats** |  |  |  |  |  |  |
| Effectors | 2092 | 2051 | 2178 | 2036 | 1928 | 1956 |
| Singleton effectors<sup>h</sup> | 124 | 162 | 177 | 180 | 137 | 139 |
| Effector with orthology from other isolates<sup>i</sup> | 653 | 574 | 535 | 390 | 385 | 411 |
| Synteny effector pairs | 890 | 890 | 1099 | 1099 | 1121 | 1121 |
| Chromosome effector pairs | 297 | 297 | 269 | 269 | 233 | 233 |
| Unpaired region effector pairs | 128 | 128 | 98 | 98 | 52 | 52 |

a. "P" refers to primary contigs.
b. "H" refers to haplotigs.
c. single copy orthologs: one orthology present in each of the 6 genomes of 3 *Pt* isolates.
d. singletons: no orthologs across all the remaining genomes of 3 *Pt* isolates.
e. synteny allele pairs: within clustered homologous regions of haplotig and primary contigs
f. chromosome allele pairs: out of clustered homologous regions, but located on primary and haplotig contigs which are corresponding to each other
g. location-umatched ortholog pairs: the locations of ortholog pairs have no corresponding relationship
h. singleton effectors: genes of a diploid organism that lack both paired alleles within an isolate and orthologs across isolates.
i. no paired alleles within the isolate but at least one ortholog from other isolates

For each isolate, we predicted about 2,000 effectors on the primary contigs (**Supplementary file 4**) and haplotigs (**Table 2; Supplementary file 4**), respectively, comprising about 8% of the total proteins in each genome, in line with previous reports for *Pt* Race1, *Pst*, and *Pca* (11, 18, 19). A majority (57-70%) of the effectors belonged to either synteny or chromosome allele pairs, and only 6-9% of the effectors were singletons (**Table 2**). As genes in allele pairs contain fully phased protein sequences and in most cases the virulence gene is homozygous at the mutation point, the phased effector sequences in pairs may provide additional layer of information for prioritizing candidates of avirulence genes.

Databases including GO, PFAM domains, interproscan, CAZymes, MEROPS, and transcription factor families were used for functional annotation (**Supplementary file 5A**). GO enrichment analysis revealed no significant over- or under-representations, implicating similar abundances of GO terms between genomes. We detected about 400 CAZymes in each isolate assembly, of which 20-23% (81-91 CAZymes) was predicted as effectors (**Supplementary file 5B**). A previous study on rust fungi suggested that CAzymes had roles in plant cell wall degradation and fungal cell wall reorganization (24). As seen in *Pgt* and *Melampsora larici-populina* (25), the subclass of CAzymes, the GH5 (cellulase and other diverse enzymatic functions) and GH17 (β-glucosidase) families were also the most abundant families in the *Pt* isolates. Detailed results based on databases of MEROPS and transcription factor families are described in **Supplementary file 5B**.

In summary, our assemblies are clearly significantly better than previously published genome assemblies, for example the *Pt* Race1 genome (14,818 scaffolds with N_50_ of 544 kb) (11) and the recently reported genomes of *Pst* and *Pca* based on PacBio long-read sequencing (N_50_ of 1.5 Mb and 268 kb, respectively) (18, 19). To date, the three *Pt* assemblies reported here exhibit the highest level of contiguity and completeness of any rust fungus genome available, providing a fundamental platform for in-depth investigation of the genome function of this destructive pathogen and, more generally, of dikaryotic fungi evolution in the absence of sexual cycle in nature.

### Heterozygosity in the haplotype phased *Pt* genomes

The availability of the phased *Pt* genomes provides an opportunity to investigate the heterozygosity of *Pt* genomes at both levels of small variants (SNPs, indels, and multiple nucleotide polymorphisms) and structure variation (SV). Small variants were identified by mapping Illumina reads to the primary contigs using CLC genomics workbench v10.1. Within each isolate, we detected 3.5, 3.6, and 3.9 heterozygous variants/kb (including 3.0, 3.1, and 3.4 SNPs/kb) for Pt64, Pt104, and Pt53, respectively. Broadly consistent with the previous reported rate of 2.6 heterozygous SNPs/kb (11, 16), the slightly higher rate in this study (3.0-3.4 SNPs/kb) may reflect a better calling rate of heterozygous genotypes related to the significantly improved continuity of our newly generated assemblies.

Three types of SVs (insertions/deletions, tandem expansions/contractions, and repeat expansions/contractions) were identified between primary contigs and haplotigs within each isolate using Assemblytics (26). The distribution of these SVs was similar across the three isolates, with average counts of 1,564, 1,033, and 412 detected for insertions/deletions, repeat expansions/contractions, and tandem expansions/contractions, respectively (**supplementary file 6**). SVs (50 bp - 100 kb) represented 7.8-8.6% of the primary contig genome size in each of the three isolates, slightly higher than the 6.4% SV components reported for *Pst* (19). While insertions/deletions and repeat expansions/contractions were more frequent than tandem expansions/contractions, large SVs (50-100 kb) were observed for deletion (two events in each isolate), tandem expansion (1 event in Pt64), and repeat contraction (4-9 events in each isolate; **supplementary file 6**). The importance of SVs in the pathogenicity of rust fungi has been highlighted by our recent identification of *AvrSr50*, which demonstrated that it was an ~200 bp insertion in the *AvrSr50* gene that resulted in the development of virulence (17). Our three *Pt* assemblies generated from PacBio data with an average read length of > 10 kb offer many advantages for the detection of SVs as compared to short read assemblies, providing a fundamental platform for in-depth investigation of SVs in relation to rust pathogenicity.

### Hybrid nature of Pt64 demonstrated by population structure

To get a snapshot of the genetic characteristics of Pt64 in the context of *Pt* population structure, we selected an additional 27 *Pt* isolates representing a wide range of pathotypes for Illumina sequencing. The derived whole genome SNPs were subjected to structure analysis, a model-based clustering method for inferring population structure and assigning individuals to populations with the estimation of admixture proportions (27, 28). Three major groups across 30 *Pt* isolates were identified and the structure plot (**Figure 1a** for structure plot; **Figure 1b** for phylogenetic tree of mating type genes from 30 *Pt* isolates) clearly showed that while the presumed parental isolates Pt53 (S365) and Pt104 (S423) belong to two distinct groups, Pt64 (S473) is an admixed race with 37% and 63% proportions belonging to Pt53 and Pt104 groups, respectively. In addition, six more isolates also displayed various admixture patterns (**Figure 1a**; S634, S646, S523, S631, S543, and S564), all of which could represent previously unidentified SH isolates. Interestingly, one of these (S543) was identified as a spontaneous greenhouse mutant derived from Pt64 (S473), with added virulence for *Lr41*. While the admixture pattern gives the initial confirmation of the hybrid nature of Pt64 from the perspective of population genetics, the observations of additional SH isolates indicate that SH likely occurs in wheat rust pathogens more frequently than previously thought, increasing the potential for this pathogen to accelerate host adaption in the absence of sexual recombination.

**Figure 1.**
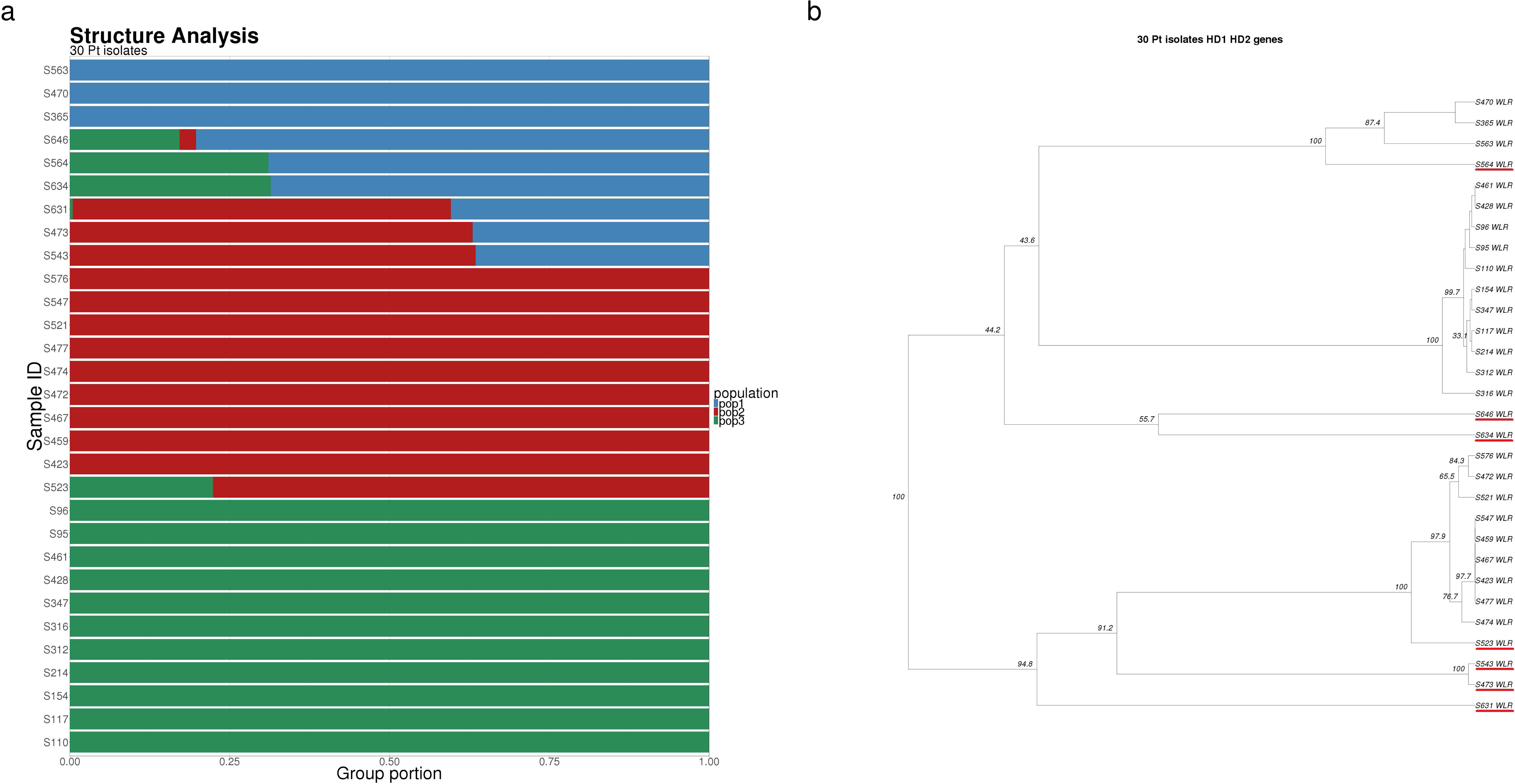
a) Structure plot of the 30 *Pt* isolates. Bars represent an isolate and the color proportion for each bar represents the posterior probability of assignment to one of the three clusters. b) Phylogenetic tree of HD genes for 30 *Pt* isolates inferred from SNPs identified for HD1 and HD2 genes. Admixed isolates being potential hybrids were underlined in red.

To further inspect the hybrid nature of Pt64, trio analysis implemented in CLC genomics workbench was also carried out for the Illumina sequencing of the three *Pt* isolates, with Pt104 and Pt53 as the presumed parental isolates and Pt64 as the progeny. This approach demonstrated that 43.5% of the small variants in Pt64 showed a biparental origin from Pt53 and Pt104. However, 42.5% of the variants had the origins undifferentiable between the parental isolates (**supplementary file 7**), which could be attributed to genetic similarity between clonal lineages and the limitations of small variants analysis.

### Pairwise alignments of phased genomes and phylogeny of single copy BUSCO orthologs

To overcome the limitations of the small variant analysis and further inspect relationships among the three *Pt* isolates, whole-genome alignments were performed with the mummer toolset in a pairwise manner using the six phased assemblies (18, 29). The generated mummer plots (**Figure 2a**) showed that the genome similarity between Pt64 and the haplomes of parental isolates (Pt53 and Pt104) is higher than that between parental haplomes, demonstrating divergence between parental isolates and convergence between the hybrid and its parental isolates, consistent with a hybrid origin of Pt64 from the genome scale.

**Figure 2.**
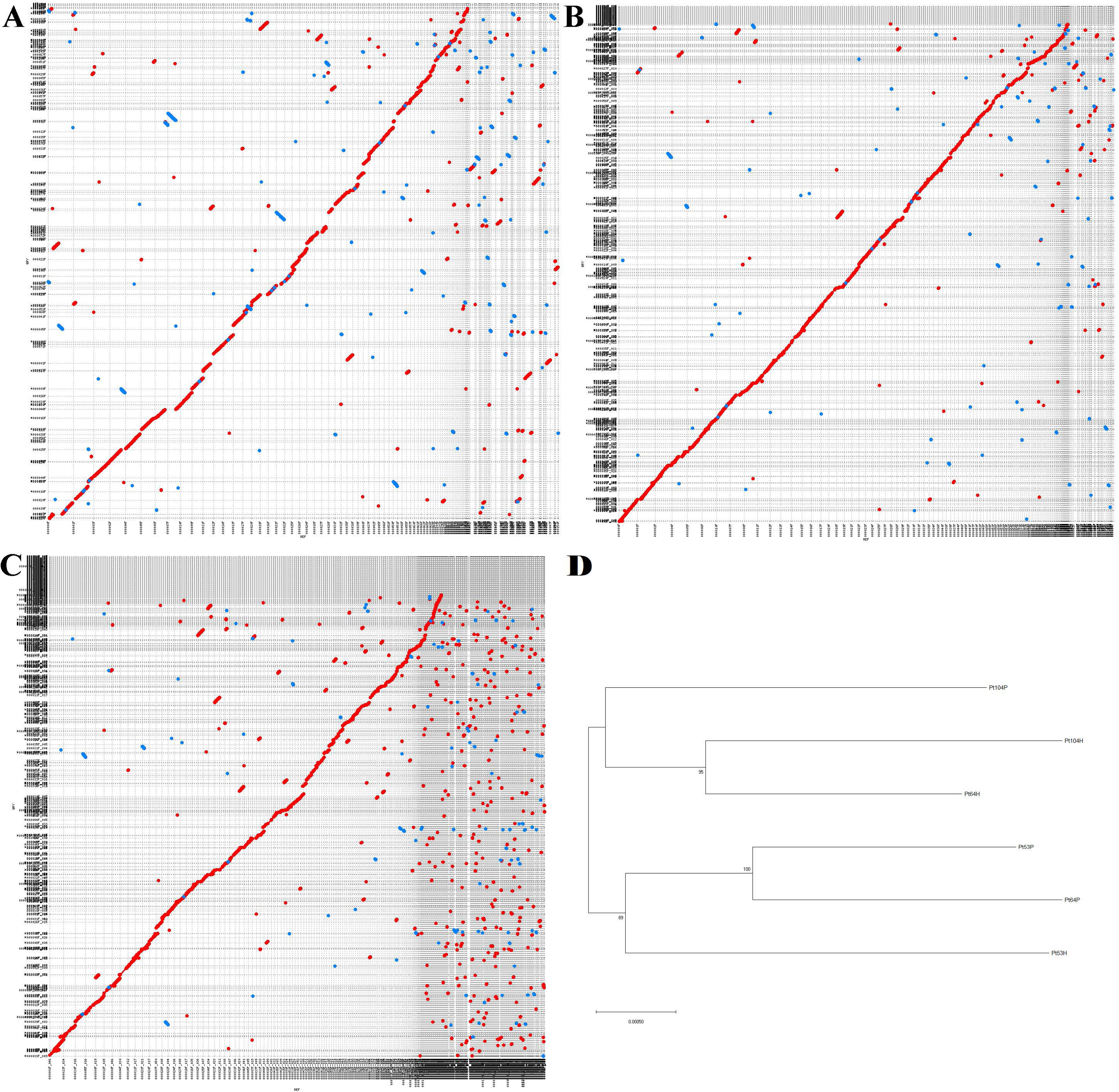
Mummer plots for the *Pt* assemblies. a) Mummer plot for alignments between Pt53 and Pt104 primary contigs. Pt53 and Pt104 contigs are labelled along horizontal and vertical axis, respectively. b) Mummer plot for alignments between Pt53 haplotigs and Pt64 primary contigs. Pt64 and Pt53 contigs are labelled along horizontal and vertical axis, respectively. c) Mummer plot for alignments between Pt104 haplotigs and Pt64 haplotigs. Pt64 and Pt104 contigs are labelled along horizontal and vertical axis, respectively. Red and blue colors indicate same and opposite strand direction. d) Phylogenetic tree of single copy BUSCO orthologs derived from the six phased genomes of three *Pt* isolates.

Gene orthology across the six phased genomes from the three isolates was also examined and 30,640 orthologous groups were identified, with each group containing 2-18 members (**Supplementary file 8**). Of these orthologs, 689 single copy BUSCO orthologs were further detected across all six genomes, the genomic locations of which are shown in **Figure 3** track III (**Supplementary file 8**). A phylogenetic tree inferred from these conserved BUSCO orthologs based on the amino acid sequences showed that the primary and haplotig genomes of Pt64 formed two distinct clusters with one cluster including primary contigs of Pt64 and the two phased assemblies of Pt53 only, and the other including haplotigs of Pt64 and the two phased genomes of Pt104 only (**Figure 2b**). This phylogeny based on amino acid sequences of functionally conserved proteins is again consistent with a hybrid origin of Pt64.

**Figure 3.**
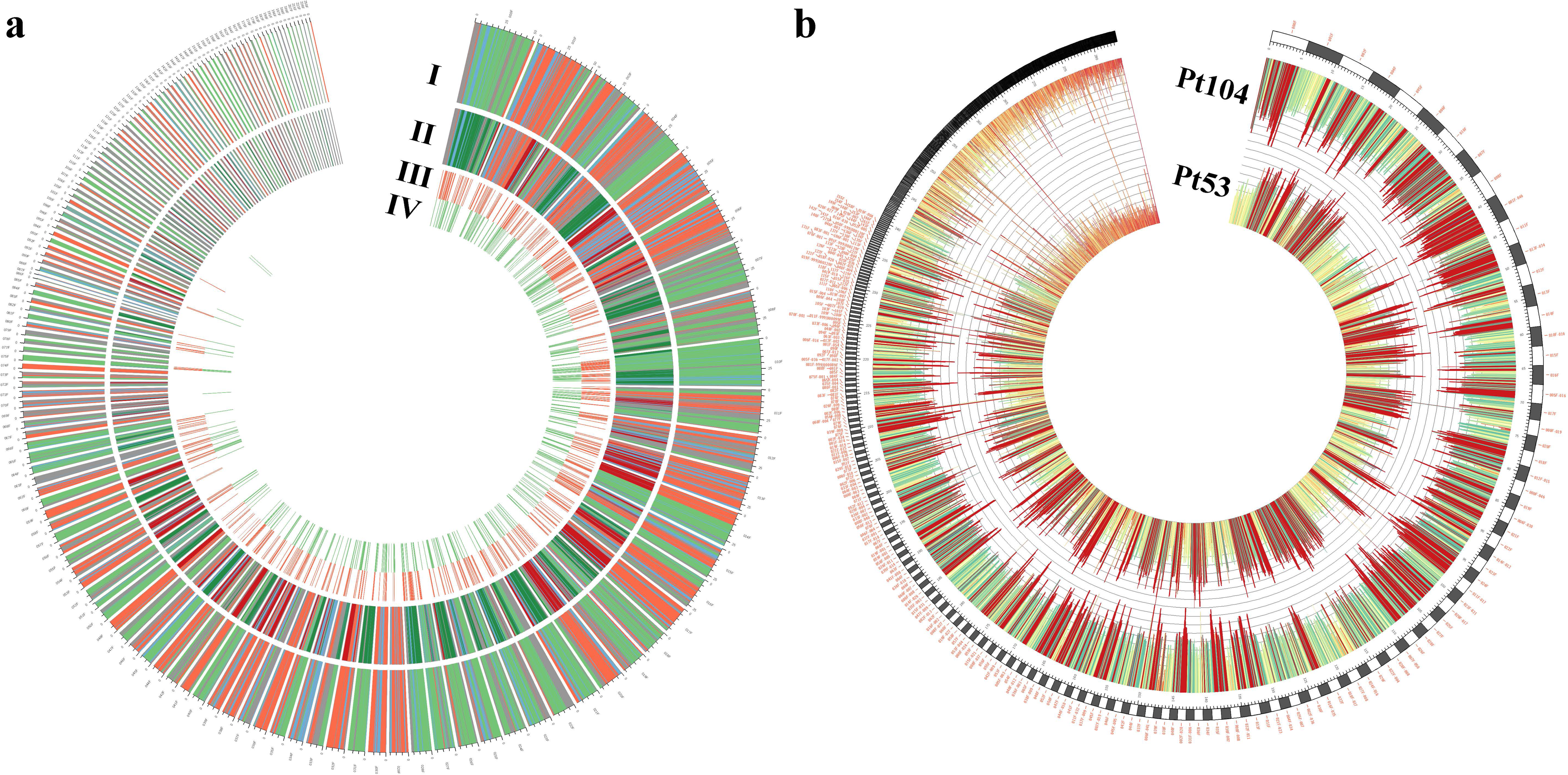
Whole genome alignments for dissecting the hybrid origin of Pt64. a) parentage of primary contigs of Pt64 and genomic locations of two gene panels. I) The primary contigs of Pt64 are dissected into 4 categories based on the alignment situations, including alignment regions unique to Pt53P and/or Pt53H (red), unique to Pt104P and/or Pt104H (green), overlapping between Pt53 and Pt104 (blue), and with no high similarity with either isolate (grey). Each tick indicates 0.1 Mb length. II) Corresponding to I, the alignment regions on Pt64 were further differentiated with phased status by primary contigs and haplotigs of Pt53 and Pt104. Pt53P and Pt53H are in orange and red, respectively. Pt104P and Pt104H are in light green and dark green, respectively. III) The genomic locations of 689 single copy BUCSO orthologs on Pt64P. IV) The genomic locations of 524 single copy hybrid genes on Pt64P. b) coverage histograms for long-read sequencing of Pt53 and Pt104 mapped against Pt64PH (concatenated contigs of both primary and haplotig contigs) individually. The colors in this diagram represent coverage values, with red being high coverage and yellow/green being low. The coverage was calculated using bedtools with 140 kb windows and 70 kb steps.

### Defining parental contributions to the hybrid Pt64 genome

To further investigate the relationship between Pt64 and its parents Pt53 and Pt104, an attempt was made to identify the parental origin of the contigs in the dikaryotic Pt64 assembly. By stringently aligning the phased parental genomes to Pt64 primary contigs and haplotigs individually (alignment length ≥ 10 kb and similarity ≥ 99.8%), 153 primary contigs (**Figure 3a**) and 412 haplotigs of Pt64 (Pt64H) were selected for parentage assignment (**Supplementary file 9**). Based on alignment status, each of the 153 primary contigs of Pt64 (Pt64P) was dissected into genomic regions belonging to one of four categories: unique to Pt53P and/or Pt53H (no match with Pt104); unique to Pt104P and/or Pt104H (no match with Pt53); overlapping between Pt53 and Pt104 (double match); and no high similarity with either isolate. While in Pt64P, 61 and 78 contigs had alignment regions dominated by Pt53 origin (maximum covering percentage of total contig length; e.g. contig 001F largely in orange in **Fig 3**) and Pt104 origin (e.g., 000F largely in green), respectively, the alignment regions assigned to Pt53 and Pt104 origins covered about 29.1% and 33.4% of the total length of the 153 primary contigs of Pt64P, respectively. For the selected 412 haplotigs of Pt64 haplomes (Pt64H), 116 and 250 haplotigs had alignment regions dominated by Pt53 and Pt104, respectively, whereas the alignment regions indicating Pt53 and Pt104 origins covered 27.3% and 41.5% of the total length of the selected 412 haplotigs, respectively. At this stage of genomic dissection, the parentage origins were differentiated only between isolates and the contribution of primary contigs and haplotigs of the same isolate was merged. By identifying the most likely parent for the majority contigs of Pt64, the results unambiguously demonstrated Pt64 is a somatic hybrid derived from Pt53 and Pt104 lineages. Interestingly, the dissection patterns of parental origin in Pt64 contigs correspond well with the reciprocal pattern of peak coverages between the parents in the coverage analysis based on PacBio sequencing of Pt104 and Pt53 mapping to concatenated contigs of Pt64 primary and haplotig contigs individually (**Figure 3b**; **Supplementary file 9)**, which further confirmed the reliability of our parentage assignment.

### Haplotype dissection reveals a mechanism more complex than simple nuclear exchange

Although SH has long been proposed for rust fungi by laboratory studies, the underlying mechanisms are unknown. To answer this question, we inspected the parentage of haplotypes without potential haplotype switching, which can represent genomic regions from a single nucleus (30). These haplotypes included haplotigs from Pt64H and haplotypes within phase blocks (a primary contig segment uniquely matched by the corresponding haplotig). Of the 412 haplotigs selected as aforementioned, 43 had biparental origins and the alignments representing Pt53 and Pt104 origins covered 39% and 20% of the total length of these haplotigs, respectively (**Figure 4a**). Within phase blocks, 57 primary contigs had biparental origins (**Figure 4b**), and the segments implicating Pt53 and Pt104 origins covered 32% and 30% of the total length of these primary contigs, respectively. These observations of biparental origins in a genomic region from a single nucleus indicate that mitotic recombination occurred in at least one nucleus of Pt64. Furthermore, for 26 phase blocks, both primary contig and haplotig within the block were found to have biparental origins (**Figure 4c**). As each haplotype within a phase block contains a genomic region from a different single nucleus, concurrent biparental origins in both haplotypes within a phase block suggests the occurrence of somatic crossover in both nuclei of Pt64. Altogether, our findings clearly demonstrate that the mechanism underlying SH in *Pt* definitely involves mitotic crossover, a mechanism more complex than simple nuclear exchange as previously thought.

**Figure 4.**
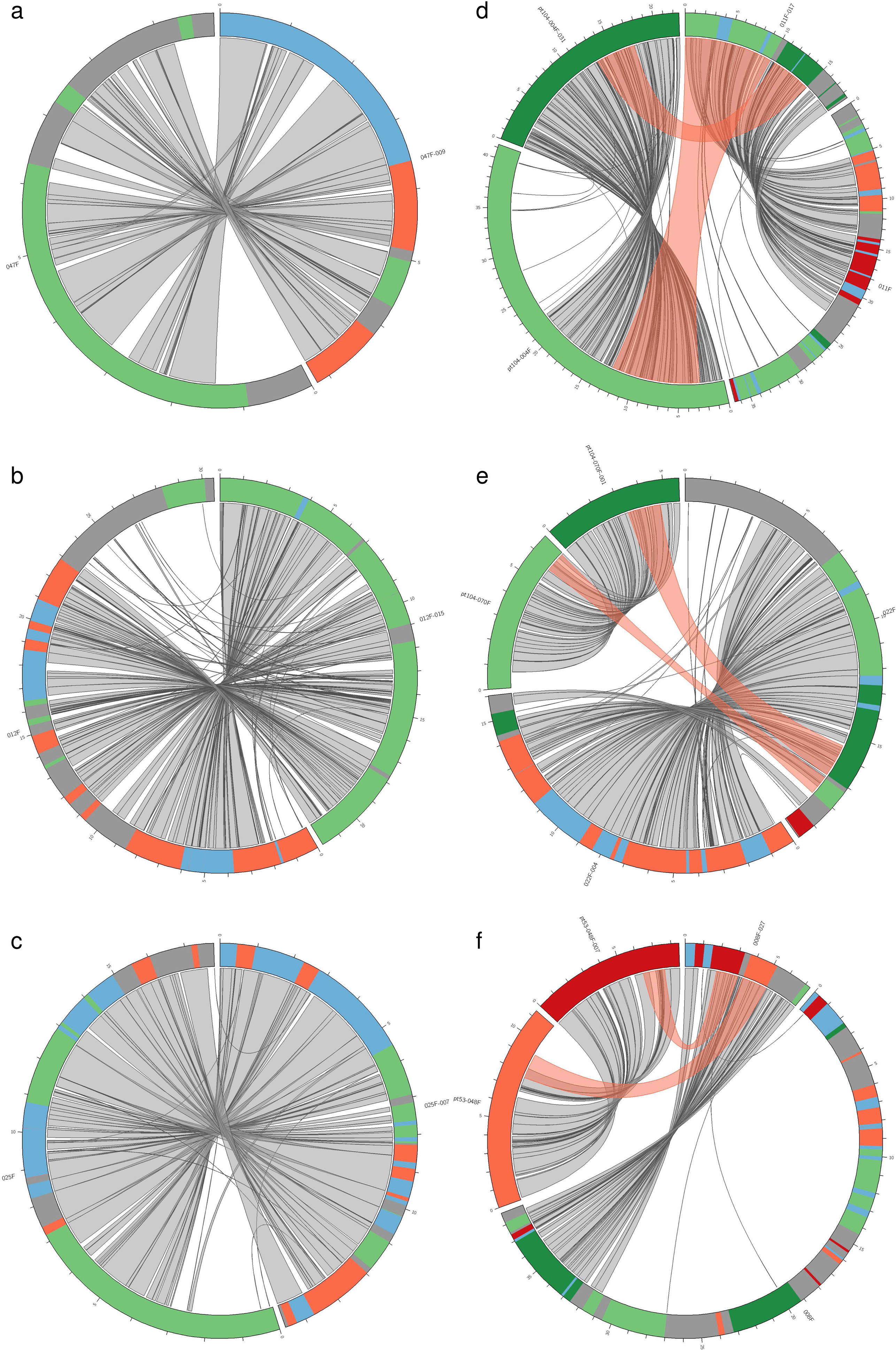
Examples of biparental origins of haplotypes implicating the occurrence of mitotic crossover and examples of complex recombination patterns showing one nucleus of Pt64 had components from both nuclei of a parental isolate. The corresponding regions between primary and haplotig contigs are shown by gray shading. Color coding for a-c: orange, green, blue, and gray regions represented for alignment regions unique to Pt53, Pt104, overlapping in Pt53 and Pt104, and no high similarity alignment, respectively. a) Pt64 haplotig 047F_009 had alignment regions with Pt53 and Pt104 origins simultaneously. b) Pt64 primary contig 012F with the segment of biparental origins within one phase block (no phase switch on these segments). c) primary contig 025F and corresponding haplotig 025F_007 of Pt64, both of which had biparental origins within the homologous matching region (no phase switch within one phase block, representing sequences from each of the 2 nuclei), demonstrating occurrence of somatic crossover in both nuclei. Color coding for d)-f): Pt53P and Pt53H are in orange and red, respectively. Pt104P and Pt104H are in light green and dark green, respectively. Blue and gray use the color coding as aforementioned. d) a Pt64 haplotig 011F_017 with alignments of both Pt104P and Pt104H origins (pink shading), and the segments of the two haplomes of Pt104 matched to Pt64 were located in one phase block (homologous matching) without phase switching (left part of fig d), indicated that haplotig 011F_017 combined sequences from two nuclei of Pt104. e) the recombination pattern observed in primary contig 022F of Pt64, demonstrating that 022F combined sequences from two nuclei of Pt104. f) the recombination pattern of Pt64 haplotig 008F_027 demonstrating that one nucleus of Pt64 had components from two nuclei of Pt53.

### Contributions of multiple parental nuclei to a single nucleus of Pt64 by mitotic crossover

The aforementioned parentage analysis can be further refined with the differentiation between primary contigs and haplotigs of Pt53 and Pt104 (Pt53P, Pt53H, Pt104P, and Pt104H), which showed that all four parental genomes contributed to the Pt64 genomes (**Figure 3a track II**; **Table 3; Supplementary file 9**). To exclude the potential impact of haplotype switching, we screened the haplotypes of Pt64 and its inherited parental regions within haplotig and phase blocks. We found that haplotig 011F_017 in Pt64 had parental origins of both Pt104P and Pt104H (**Figure 4d**) and the matched parental regions on Pt104 primary contigs and haplotigs were within a phase block, indicating that a single nucleus of Pt64 had parental origins from two nuclei of Pt104. Combined with our previous conclusion that both nuclei of Pt64 had origins from Pt53 and Pt104, we reached the further conclusion that at least one nucleus of Pt64 had origins from three parental nuclei (two nuclei of Pt104 and one nucleus of Pt53). Similar conclusions can be drawn for the recombination pattern observed in primary contig 022F within a phase block (**Figure 4e**) and for the haplotig 008F_027 of Pt64 (**Figure 4f**; origins from two nuclei of Pt53 and one nucleus of Pt104). Therefore, our observations demonstrated that at least one nucleus of Pt64 had genetic origins from three parental nuclei and we postulate that both nuclei of Pt64 may have genetic origins from multiple parental nuclei. This finding and our postulation are supported by laboratory studies of SH in *Pgt* suggesting that during hyphal fusion, more than two nuclei might come together and exchange chromosomes (31), and the study of germinating urediniospores of *Pt* reporting germ tube fusion bodies (GFBs) containing four nuclei with bridge-like connections between nuclei, a structure similar to the chromosomal bridges involved in chromosomal alterations (12).

**Table 3.** Statistics for alignment of parental genomes with Pt64 genomes (length >10 kb and identity > 99.8%)

|  | **53PH<sup>a</sup>** | **53P<sup>b</sup>** | **53H<sup>c</sup>** | **104PH** | **104P** | **104H** |
| --- | --- | --- | --- | --- | --- | --- |
| **alignment to 64P** |  |  |  |  |  |  |
| **Contig number of dominated origins<sup>d</sup>** | 61<sup>f</sup> | 22<sup>g</sup> | 21<sup>g</sup> | 78<sup>f</sup> | 24<sup>g</sup> | 30<sup>g</sup> |
| **Total length %<sup>e</sup>** | 29.1 | 14.3 | 13.5 | 33.4 | 13.6 | 18.6 |
| **alignment to 64H** |  |  |  |  |  |  |
| **Contig number of dominated origins** | 116 | 92 | 25 | 250 | 110 | 135 |
| **Total length %** | 27.3 | 15.6 | 11.5 | 41.5 | 18.4 | 21.8 |

a: "PH" refers to primary and haplotig contigs.
b: "P" refers to primary contigs.
c: "H" refers to haplotigs.
d: the total number of Pt64P contigs dominated by alignment regions unique to a parental isolate assembly (e.g., 53P, or 53H, or 53PH). Alignment regions unique to a parental isolate showing the maximum covering percentage of total contig length was considered as the dominated origin.
e: the total length of the alignment regions unique to a parental isolate assembly versus the total length of the 153 primary contigs of Pt64P.
f: The contig number was derived from differentiation between Pt53 and Pt104 only, regardless of primary contigs and haplotigs.
g: The contig number was derived from differentiation between Pt53P, Pt53H, Pt104P and Pt104H simultaneously.

Although laboratory studies of *Pgt* (17) and *Pst* (31) have provided evidence of SH under artificial laboratory conditions, with the former implicating somatic exchange within an isolate resulting in loss of *AvrSr50* and the latter providing evidence of new virulence combinations derived from parental isolates, our study is the first to confirm SH in nature and to reveal the complex process of mitotic recombination in SH. While simple nuclear exchange or reassortment produces a new pathotype via the replacement of entire sets of genes from another pathotype, mitotic recombination can introduce genetic changes nearly anywhere in the genome (32). By overcoming clonal interference, combining beneficial mutations arising from various genetic backgrounds, and accelerating host adaption in a way similar to sexual recombination, mitotic recombination in rust pathogens may occur in nature more frequently and hence pose a much greater threat to plant production than was previously thought. This is supported by the detection of additional six potential SHs in our population sequencing and the fact that Pt64 rendered several hybrid wheat cultivars susceptible to leaf rust in the absence of sexual recombination.

### Identification of mating type genes in the three *Pt* isolates

Recently it has been suggested that the maintenance of mating type genes in dikaryotic fungi could have more profound meaning in multiple aspects of life style than just regulating sexual reproduction (33). Given the crucial functions and lack of understanding of these mating type genes in wheat rust fungi, we investigated the mating type genes across the phased genomes of the three *Pt* isolates. The orthologs of all the mating type genes reported in *Pt* Race 1, including Mfa genes encoding putative pheromone precursors, STE genes encoding pheromone receptors, and HD genes (HD1 and HD2) encoding homeodomain-containing transcription factors (11), were successfully identified in our *Pt* genomes (**Table 4**). The close proximity reminiscent of the P/R organization found in several basidiomycetes including *Pt* Race 1 was present for the PtSTE3.2 and Ptmfa2 genes, but not for STE3.1, STE3.3 or mfa1/3 genes. HD genes were detected in both genomes of each isolate, located on contigs different to those harboring STE and mfa genes. While the genes encoding pheromone and pheromone receptor system (designated as *a* locus in *U. maydis*) mediate recognition of mating partners and cell fusion, the HD genes (*b* locus) regulate post-mating behavior by forming heterodimers that act as a transcription factor (34, 35). Because variations in the *a* locus were limited (**Supplementary file 10)** and the *b* locus has been implicated to be more involved in pathogenicity development than the *a* locus, our subsequent analysis focused on *b* locus HD genes.

**Table 4.** Mating type genes identified in the phased genomes of 3 Pt isolates.

| **Gene** | **Pt64 primary contigs** | **Pt64 Haplotigs** | **Pt104 primary contigs** | **Pt104 Haplotigs** | **Pt53 primary contigs** | **Pt53 Haplotigs** |
| --- | --- | --- | --- | --- | --- | --- |
| bW2-HD1<sup>a</sup> | GN64P176_007042 | GN64H843_007650 | GN104ID162_012070 | GN104ID713_013572 | GN53ID155_001018 | GN53ID531_000798 |
| bE2-HD2<sup>a</sup> | GN64P176_007041 | GN64H843_007649 | GN104ID162_012071 | GN104ID713_013573 | GN53ID155_001019 | GN53ID531_000799 |
| STE3.1<sup>a</sup> | GN64P176_004548 | GN64H843_005035 | GN104ID162_010826 | GN104ID713_012452 | GN53ID155_006664 | GN53ID531_006779 |
| STE3.2<sup>a</sup> | GN64P176_025004 | none | GN104ID162_021613 | none | GN53ID155_015260 | none |
| STE3.3<sup>a</sup> | GN64P176_026757 | none | GN104ID162_026103 | none | GN53ID155_023197 | none |
| mfa1/3<sup>a</sup> | 000072F:105505-105685 +<br>000072F:140168-139988 - | none | 000083F:25069-25249 +<br>000083F:59733-59553 - | none | 000082F 68604-68784 +;<br>000082F 103173-102993<br>- | none |
| Mfa2<sup>a</sup> | 000077F:271814-271715 - | none | 000049F 533972-534071 + | none | 000027F:1668070-<br>1668169 + | none |

a: the orthologs of bW2-HD1, bE2-HD2, STE3.1, STE3.2, STE3.3 in *Pt* Race 1 are PTTG\_27730, PTTG\_03697, PTTG\_28830, PTTG\_09751, and PTTG\_09693, respectively.

### High diversity within specificity region at *b* loci is associated with admixed *Pt* isolates

The orthologs of bE2-HD2 (PTTG_27730 from *Pt* Race 1) and bW2-HD1 (PTTG_03697) on both primary and haplotig contigs in each of the three isolates were found and each pair of HD1 and HD2 genes was divergently transcribed, encoding polypeptides of 612-679 and 373-450 amino acids, respectively. Alignment of protein sequences revealed that the HD1 and HD2 protein sequences of Pt64P and Pt64H are uniquely identical to those from haplotigs of Pt104 and Pt53, respectively. Interestingly, such a hybrid pattern in Pt64 (uniquely identical protein sequences with one counterpart of each parent) was also present in 524 single copy orthologs (**Figure 3a track IV**), which along with the aforementioned phylogeny of BUSCO orthologs convergently manifested the hybrid nature of Pt64 at protein sequence level.

Previous studies on mating type genes have demonstrated that between different mating types, self and non-self-recognition is governed by the dimerization of HD proteins (e.g. bE1 with bW2; **Figure 5a**;), which is mediated by the N terminus of HD proteins in *U. maydis*, the most different region on the *b* locus (**Figure 5a**) (36, 37). Within this highly variable region related to the specificity of mating type, we observed the breaking down of sequence similarity between Pt64P and Pt64H contrasting with the high similarity maintained between Pt64P and Pt104H as well as between Pt64H and Pt53H. These results clearly demonstrate that the 100 kb regions harboring the *b* locus (**Figure 5b**) in Pt64P and Pt64H have distinct origins closest to the counterparts in Pt53 and Pt104 haplotigs, respectively, again underscoring the hybrid nature of Pt64.

**Figure 5.**
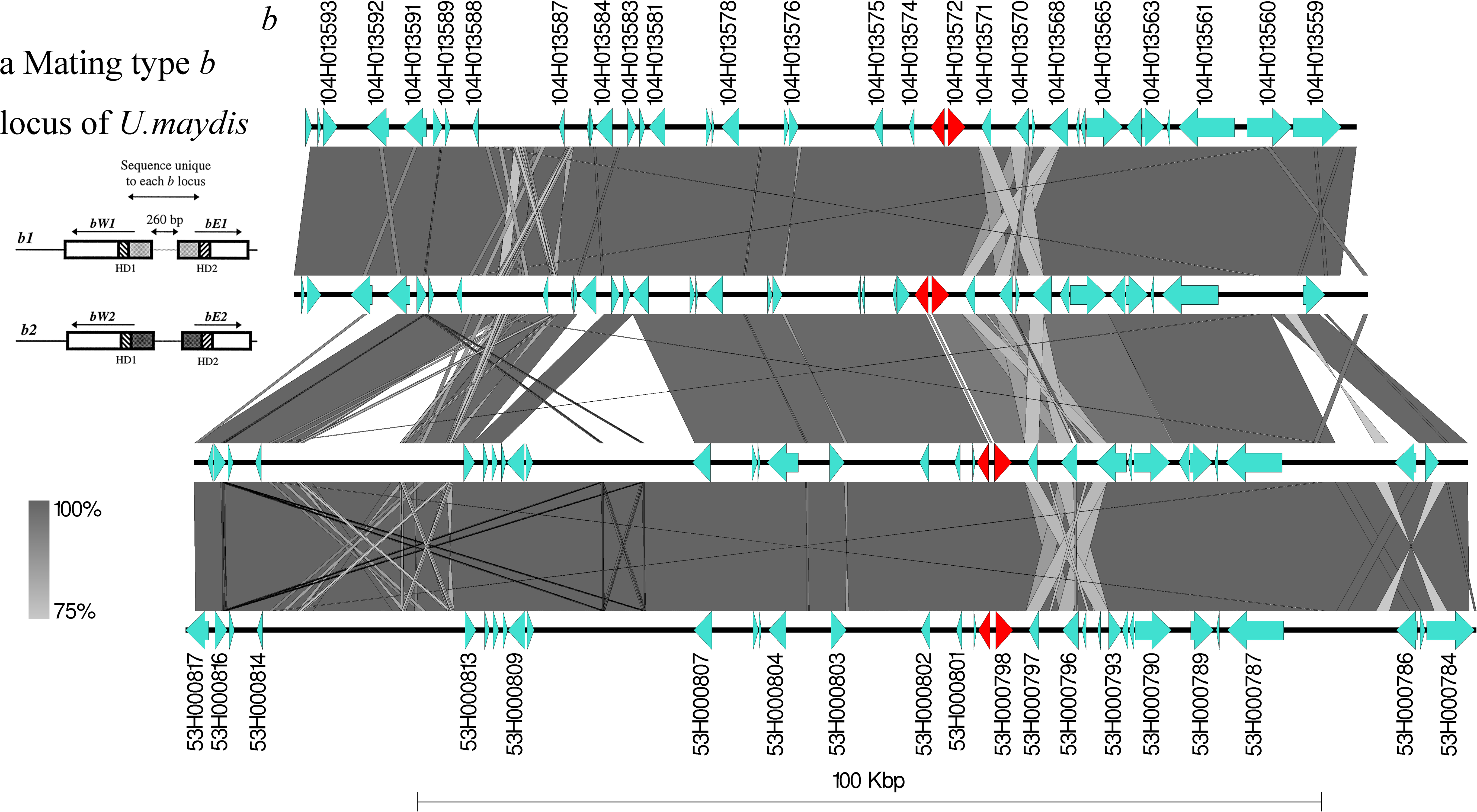
a) Molecular structure and organization of the *b* mating-type locus of *U. maydis*. The b1 and b2 loci are illustrated, and the variable DNA sequence specific to each allele is highlighted by light and dark grey, respectively. Straited boxes indicate the homeodomain. Compatible combinations are bE1 + bW2 and bE2 + bW1. Figure adapted from Casselton *et al*. (1998). b) Synteny analysis of ~100 kb region harbouring HD1 and HD2 genes with Pt104H, Pt64P, Pt64H, and Pt53H assemblies from the top to the bottom. Gene models are represented as arrows and labeled with their locus tag for Pt104H and Pt53H. HD1 and HD2 genes are highlighted in red. Vertical gray shading illustrates the blastn identity between sequences, according to the scale shown in the right bottom corner next to the sequence scale bar.

We further asked whether the high variation within the specificity region on the *b* loci from each haplome of Pt64 arose by chance, or could it be intrinsically connected with the SH? To get a clue, a phylogenetic tree based on SNPs within the HD1 and HD2 genes was constructed, which demonstrated that all seven admixed races identified by STRUCTURE analysis occupied branches distinct from non-admixed members within the same clade (**Figure 1b**). Most prominently, the admixed isolates, S646 and S634, along with S543 and S473 (Pt64), formed two distinct sister groups within each of the two major clusters. Although accurate phased sequences were unavailable for the additional 27 isolates due to the limitation of short reads, the distinct pattern shown by admixed isolates suggests that HD genes in nature-derived hybrids may possess features distinct from non-admixed members, and that a high level of genetic variation in specificity-related regions between HD genes seen in Pt64 could be intrinsically associated with the SH event. One possible explanation could be that such a high diversity may be necessary for the successful heterodimerization of the HD genes in the SH process, which generates active transcription factors leading to a new developmental pathway (37). This pathway may involve biological processes crucial for a successful SH, such as DNA damage repair via homologous recombination, which has been suggested as the most plausible reason for maintaining the mating type genes in dikaryotic fungi without sexual reproduction (33).

## Conclusions

Several mechanisms, including exotic incursions, single-step mutation, and SH are postulated as causal factors of virulence changes in populations of asexually reproducing cereal rust pathogens in Australia (2). While comparative genomics mainly focuses on DNA mutations, few studies have specifically characterized molecular events associated with SH and none have revealed how this process happens in nature. The most significant implication of our study is that, for the first time, the genomic characteristics of a naturally occurring SH in a rust pathogen have been unraveled and the underlying mechanism involving complex pattern of mitotic crossover has been uncovered.

In brief, using third generation long-read sequencing, the first phased dikaryotic genomes of *Pt* for a putative hybrid isolate and the two parental isolates were generated. The genetic dissection based on the high-quality phased genomes including whole-genome alignments, phylogenetic and syntenic analyses along with population sequencing convergently highlighted the hybrid feature of Pt64 genomes. Parentage analysis of the three phased *Pt* genomes within haplotig and phase block regions further demonstrated the role of mitotic crossover in the derivation of both nuclei of this SH isolate. Furthermore, within HD mating type genes, the distinct specificity regions in Pt64 and the distinct phylogenetic pattern of the remaining admixed isolates suggested a high level of genetic variation in specificity-related regions on *b* locus intrinsically associated with the SH. Our study has not only provided a fundamental platform for investigating genomic variation and function underlying virulence development in one of the most devastating pathogens of wheat, but has also provided an in-depth understanding of the mechanisms of naturally occurring SH. This mechanism can be exploited by any dikaryotic pathogen in nature to accelerate virulence evolution in the absence of sexual recombination, and understanding this process will undoubtedly facilitate more sustainable pathogen control. Moreover, this study has provided a new paradigm to investigate naturally occurring SH in dikaryotic pathogen with the obligate biotrophic life style, a feature that hampers the ability to culture the fungus *in vitro* and hence limits downstream biological and genetic studies based on culturing. In future, combining Hi-C data with our current genome assemblies may yield fully phased chromosome-length scaffolds to further improve our understanding of somatic exchange and virulence evolution in dikaryotic pathogen.

## Methods

### Puccinia triticina isolates

Pathotype 104-2,3,(6),(7),11 (isolate 840045=S423) was first detected in Victoria in 1984 and concluded to be an exotic incursion based on the observations of pathogenic and isozymic differences to isolates of *P. triticina* endemic at that time (38). A second pathotype, also regarded as having an exotic origin, pt 53-1,(6),(7),10,11 (isolate 810043=S365), was first detected in Australia in 1984 after being initially detected in New Zealand in 1981. Pathotype 64-(6),(7),(10),11 (isolate 900053=S473) was first detected in northern New South Wales in 1990. This pathotype was believed to have arisen via SH between pt 53-1,(6),(7),10,11 and 104-2,3,(6),(7),11 based on comparative studies of pathogenicity, isozymes, and RAPD fingerprinting (4). In addition, 27 additional *Pt* isolates with a range of avirulence/virulence attributes (“pathotypes”) were also included for phylogenetic analysis of population structure (**Figure 1a**) and HD mating type genes (**Figure 1b**).

### Plant inoculation

To ensure purity of each isolate for PacBio sequencing, a single pustule was selected from a region of low-density infection and propagated on wheat plants of the susceptible variety Morocco prior to DNA preparation. The identity and purity of each isolate were checked by pathogenicity tests with a set of host differentials at each cycle of inoculum increase and also using urediniospores subsampled from those used for DNA extraction. For rust infection, plants were grown at high density (~25 seeds per 12 cm pot with compost as growth media) to the one leaf stage (~7 days) in a greenhouse microclimate set at 18–25°C temperature and with natural day light. Plants were inoculated as previously described ^42^. For DNA isolation, mature spores were collected, dried and stored at −80°C.

### DNA extraction and genomic DNA sequencing

DNA was extracted from urediniospores as previously described (39) and PacBio sequencing was performed at the Australian Genome Research Facility Ltd (Adelaide, Australia). For library preparation, the 20 kb BluePippin kit (PacBio) was used and DNA libraries were sequenced on a PacBio Sequel System with Sequel Sequencing chemistry 2.1. We sequenced 3-4 SMRT cells per sample and each SMRT cell had a capacity of 5-10 Gb per cell. For Illumina short-read sequencing, TruSeq library of DNA samples from the same *Pt* pathotypes were constructed with a 150 bp paired end and sequenced on a HiSeq 2000 instrument at Novogene (Hong Kong, China).

### Genome assembly and curation

The integrated pipeline of FALCON and FALCON-Unzip (v4.1.0) was used for genome assembly (30). Read length cutoffs were computed by FALCON based on the seed coverage and expected genome size. After assembly by Falcon, FALCON-Unzip was used to phase haplotypes and to generate consensus sequences for both primary contigs and haplotigs (the latter being the phased haplotype blocks associated with the former), which were then subjected to further correction using Quiver implemented in SMRT (v4.0.0). Blastn searches against the NCBI nucleotide reference database were used to check potential noneukaryotic contamination and none of the contigs were found to have predominant noneukaryotic sequences as best BLAST hits at any given position. These assemblies were further curated and polished by removing low quality contigs and reassigning primary contigs without haplotigs showing a significant match with another primary contig. We performed three manual curation steps using the following criteria for removing low quality contigs or reassigning primary contigs: (1) contigs with extreme low or high coverage (coverage < 10 or > 2000 fold) were removed; (2) contigs smaller than 100 kb and > 20% of the contigs showing no consensus call marked by Quiver (lowercase) were removed; and (3) primary contigs without haplotigs showing significant match (> 85% best match coverage) with another primary contig were reassigned to haplotigs (40). To evaluate assembly completeness, the software BUSCO (v3.0) (41) was used for comparison with the fungal lineage set of orthologs (basidiomycota_odb9), which consisted of 1,335 basidiomycete conserved orthologs. Telomeres were identified by the presence of at least 10 repeats of CCCTAA or TTAGGG within 200 bp of the end of a contig as previously described (18).

### SNP associated analysis and inter-haplotype structure variation

CLC Genomics Workbench (v10.1.1) was used for SNP detection. In brief, trimmed DNA reads were first mapped to the annotated primary contigs of Pt64, which were then subjected to local realignments. Fixed ploidy variant detection (ploidy = 2) was performed, ignoring non-specific matches with parameters including variant probability 90%, minimum coverage of 10, minimum variant count 2, and minimum frequency 20%. After filtering, SNP calls were subjected to trio analysis implemented in CLC Genomics Workbench, with Pt53 and Pt104 as the parental isolates, and Pt63 as the progeny. This tool reported the potential origin of a variant allele, including inherited from father, from mother, from either parent, from both, recessive, and *de novo*.

For identification of inter-haplotype structure variation (SV) within each isolate, we first used mummer3 (29) to align haplotigs to their corresponding aligned regions in primary contigs. The output file (delta file) from mummer3 was then fed into Assemblytics (26) for detecting SV between haplotypes.

Potential admixture and population structure in 30 *Pt* isolates including Pt64, Pt104, and Pt53 were investigated. Quality-trimmed Illumina reads of these 30 isolates were mapped against *Pt* Race 1 genome using bwa mem method and SNPs were detected and filtered using GATK v3.8.0, which were then fed into the software fastStructure with K values ranging between 1-8 (42). The SNPs identified in HD1 and HD2 genes were further extracted to construct a phylogenetic tree using Poppr (43).

### Parentage analysis by whole-genome alignments and coverage analysis between *Pt* isolates

Pairwise whole-genome alignments between six phased genomes of the three *Pt* isolates were performed using mummer toolset (nucmer –mum, delta-filter -l 10000 and mummerplot) (29) (18) to investigate the relationships of the three *Pt* isolates (**Figure 2**). The alignments generated were further selected for high similarity (≥ 99.8%). This optimized threshold was able to both differentiate parental isolate assemblies and to maximize the total percentages of Pt64 assemblies that parental assemblies can be aligned. Based on the alignment status, the contigs of each Pt64 assembly were further dissected into four categories, *viz*. alignment regions unique to Pt53P and/or Pt53H, unique to Pt104P and/or Pt104H, overlapping between Pt53 and Pt104, and with no high similarity with either isolate. These dissections were displayed in a circular plot using Circos (44). We also performed a detailed coverage sequence depth analysis for the three *Pt* isolates. The long-read sequencing data of Pt53 and Pt104 were mapped against Pt64PH (concatenated contigs of both primary and haplotig contigs) individually and coverage was calculated using bedtools with 140 kb windows and 70 kb steps. The mean base sequence depth was visualized in a circular histogram plot using Circos (44).

### RNA isolation and sequencing

Infected leaves were collected at 1, 2, 5, and 7 days after inoculation and immediately frozen in liquid nitrogen. Samples were ground to a fine powder in liquid nitrogen and total RNA was isolated with the isolate II RNA Mini Kit (Bioline). After DNase treatment (Promega), RNA was further purified by on-column DNase treatment and the quality was assessed using the Bioanalyzer 2100. For library preparation, around 10 μg of total RNA was processed with the mRNA-Seq Sample Preparation kit (Illumina), which was then sequenced on the Illumina HiSeq2500 platform (125 bp paired-end reads).

### Transcriptome assembly and genome annotation

For Pt64 transcriptome assembly, quality trimmed reads were first aligned to the primary contigs of Pt64 by using the CLC module large gap read mapping (default parameters) and mapped reads were extracted as fungal specific reads. The extracted reads were then used as input to build *de novo* transcriptome assembly using Trinity (v2.1.1) (45). Separately, Trinity was also used to build genome-guided transcriptome assembly with the RNA sequencing bam file generated from the CLC. These transcript models along with expressed sequence tag (EST) sequences from various life cycle stage of *Pt* (22) were then used as transcript evidence for genome annotation. Funannotate (v0.7.2; https://github.com/nextgenusfs/funannotate) was used for *Pt* genome annotation, which is a pipeline specifically developed for fungi by integrating a wide range of tools such as AUGUSTUS, GeneMark, BUSCO, BRAKER1, EVidence Modeler, GAG, tbl2asn, tRNAScan-SE, RepeatModeler, RepeatMasker, Exonerate, and GMAP. Briefly, the workflow of this pipeline includes repeat identification using RepeatModeler (v1.0.8) and soft masking using RepeatMasker (v4.0.6; http://www.repeatmasker.org/), alignment of protein evidence to the genomes using TBLASTN and exoneratet (v2.2.0) (46), alignment of transcript evidence using GMAP (47), *ab initio* gene prediction using AUGUSTUS (v3.2.1) and GeneMark-ET (v4.33) trained by BRAKER1 (48), tRNAs prediction using tRNAscan-SE (v1.3.1) (49), generating consensus protein coding gene models using EvidenceModeler (v1.1.1) (50), and final clean of removing low quality gene models. For each isolate, primary contigs and haplotigs were annotated independently. RNA-seq bam file, *Pt* Race 1 protein fasta file, and transcript models along with downloaded ESTs (for gene model training, protein evidence, and transcript evidence, respectively) were fed into the Funannotate pipeline for a comprehensive annotation.

### Secretome prediction and functional annotation

Proteins predicted to have a signal peptide with no transmembrane segment and no target location to mitochondria were identified as effector candidates. SignalP v3.0 (51), TMHMM v2.0 (52), and TargetP v1.1 (53) were used for the prediction of signal peptide, transmembrane domain, and subcellular location, respectively. Following the gene prediction module as aforementioned, functional annotation to the protein-coding genes was carried out by Funannotate using curated databases including UniProt (54), Pfam domains (55), CAZYmes (56), proteases (MEROPS) (57), and InterProScan (58).

### Protein orthology analysis and classification of allele pair status

Proteinortho v5.16 was used for searching genes on primary contigs with an ortholog present on the corresponding haplotig in synteny mode, which used the relative order of genes (synteny information) as an additional criterion to disentangle complex co-orthology relationships (59). To identify homologous haplotig and primary contig regions (phase blocks), NUCmer was used for alignments between primary and haplotig sequences and the alignment coordinates were then used for scanning aligned blocks along the primary contigs and chaining the aligned haplotig blocks located within 15 kb (synteny blocks). The ortholog pairs detected by Proteinortho were then further classified based on their location in relation to synteny blocks. Those that fell into clustered homologous regions of haplotig and primary contig (synteny blocks) were defined as synteny paired alleles, those located on a primary contig and a corresponding haplotig but out of synteny blocks were defined as chromosome paired alleles, and those remaining that did not meet either of these criteria were defined as location-unmatched pairs.

### Comparison between annotated genomes and identification of mating type genes

Funannotate was also used to compare genome annotations across the six phased genomes of the three isolates. Proteinortho was implemented in this comparison to identify orthologs across all six genomes, and genes lacking both paired alleles within the isolate and orthologs across the isolates were defined as singletons. Single copy orthologs were also identified in the comparison and single copy BUSCO orthologs were then used to infer the phylogenetic tree. For mating type gene analysis, mfa genes encoding putative pheromone precursors were searched through the whole genomes against *Pt* Race 1 mfa protein sequences using exonerate with protein2genome mode (11). The STE genes encoding pheromone receptors, and HD1 and HD2 genes in our assemblies, were detected through ortholog identification of STE, HD1, and HD2 genes of *Pt* Race 1. The synteny analysis of the 100 kb fragments harbouring HD gene between *Pt* isolates was performed using Easyfig (60).

### Availability of data and materials

All raw sequence reads generated and used in this study will be available in the NCBI BioProject PRJNA343337. Supplemental data files, genome assemblies and annotations will be available on our manuscript’s dropbox page. https://www.dropbox.com/sh/ch5bf3h9fqrf4qm/AADXpH15qcEpSWW8QXv-sRxka?dl=0

## Acknowledgments

This work was funded by the Australian Grains Research Corporation (GRDC; 9175448) and USDA CSREES (2008-35600-04693). This research was undertaken as part of a long running program on national cereal rust surveillance. The program has been hosted by the University of Sydney since 1921, and since 1990 co-funded by the Australian Grains Research and Development Corporation, a statutory corporation founded in 1990 under the Primary Industries Research and Development Act 1989 and principally funded by a grower levy and Australian Government contributions. These sources had no further role in study design, data collection and analysis, decision to publish, or preparation of the article. We would like to acknowledge the Sydney Informatics Hub at the University of Sydney for providing access to the CLC Genomics Workbench (Core Research Facilities User Access Scheme) and the High Performance Computing cluster, Artemis. Particularly, we would like to thank Bernard Kirby, Leigh Burchat, Dr Stephen Kolmann, Dr Rosemarie Sadsad, Dr Cali Willet and Tracy Chew for technical support. We would also like to acknowledge Pawsey Supercomputing Centre for providing access to High Performance Computing cluster Galaxy, Dr. Benjamin Schwessinger at the Australian National University for discussion of Falcon and Falcon-Unzip pipeline, Dr. Michael Roach at The Australian Wine Research Institute for discussion of the tool Purge Haplotigs, and Dr Ming Zhao at Arizona State University for providing access to Computing cluster GEARS, an enerGy-Efficient big-datA Research System sponsored by the United States National Science Foundation award CNS-1629888.

## Authors’ Contributions

JQW analysed the data and wrote the manuscript; CD extracted high molecular DNA required for PacBio sequencing; LS contributed to data analysis and prepared the figures; CD, LS, CAC, and RFP contributed to the manuscript; RFP designed the experiment and supervised the work. All authors read and approved the final manuscript.

---

**Supplementary file 1** Evaluation of genome completeness and repetitive DNA.

**Supplementary file 2** Annotation table for primary and haplotig assemblies for each *Pt* isolate.

**Supplementary file 3** Allele pair status for each *Pt* isolate.

**Supplementary file 4** Predicted effectors in primary and haplotig assemblies for each *Pt* isolate.

**Supplementary file 5** Functional annotations of GO terms, Interproscan, Cazyme, Merops, pfam, and transcription factors for the 3 *Pt* isolates.

**Supplementary file 6** Structure variants analysis between primary and haplotig assemblies for each *Pt* isolate.

**Supplementary file 7** Trio analysis using Pt64 as the progeny, Pt53 and Pt104 as parents.

**Supplementary file 8** Ortholog clusters of the 6 genomes of the 3 *Pt* isolate, single copy orthologs across 6 genomes, and single copy BUSCO orthologs for inferring phylogenetic tree.

**Supplementary file 9** Whole-genome alignments with stringent high similarity criteria for Pt64P and Pt64H and coverage analysis for long-read sequencing of Pt53 and Pt104 mapping to Pt64 primary and haplotig contigs.

**Supplementary file 10** Details of mating type genes on *a* locus (Mfa genes encoding putative pheromone precursors and STE genes encoding pheromone receptors,); alignment figure of STE3.1; phylogenetic tree for HD1 and HD2 genes of the phased genomes of 3 wheat leaf rust isolates.

